# Genome survey sequencing of wild cotton (*Gossypium robinsonii*) reveals insights into proteomic responses of pollen to extreme heat

**DOI:** 10.1101/2021.08.27.457898

**Authors:** Farhad Masoomi-Aladizgeh, Karthik Shantharam Kamath, Paul A. Haynes, Brian J. Atwell

## Abstract

Heat stress specifically affects fertility by impairing pollen viability but cotton wild relatives successfully reproduce in hot savannas where they evolved. An Australian heat-tolerant cotton (*Gossypium robinsonii*) was exposed to heat events during pollen development, then mature pollen was subjected to deep proteomic analysis using 57,023 predicted genes from a genomic database we assembled for the same species. Three stages of pollen development, including tetrads, uninucleate and binucleate microspores were exposed to 36°C or 40°C for 5 d and the resulting mature pollen was collected at anthesis (p-TE, p-UN and p-BN, respectively). Using SWATH-MS proteomic analysis, 2,704 proteins were identified and quantified across all pollen samples analyzed. Proteins predominantly decreased in abundance at all stages in response to heat, particularly after exposure of tetrads to 40°C. Functional enrichment analyses demonstrated that extreme heat increased the abundance of proteins that contributed to increased mRNA splicing via spliceosome, initiation of cytoplasmic translation and protein refolding in p-TE40. However, other functional categories that contributed to intercellular transport were inhibited in p-TE40, linked potentially to Rab proteins. We ascribe the resilience of reproductive processes in *G. robinsonii* at temperatures up to 40°C, relative to commercial cotton, to a targeted reduction in protein transport.

## INTRODUCTION

Attempts to develop crops resilient to heatwaves are increasingly important as we experience changes in the Earth’s climate, particularly global average temperatures which are predicted to increase up to 2°C during 21^st^ century (IPCC, 2021). Extreme temperatures are considered to be detrimental to vegetative and reproductive processes in crops, ultimately diminishing yield. The global yield of major crops is likely to diminish by approximately 3 to 7% for each degree-Celsius increase in average temperature (Zhao et al., 2017). Detailing the cellular responses to heat stress at precise developmental stages of germ cell lines will help us define the genetic markers required for molecular plant breeding of thermotolerant crops.

For plants to reproduce sexually, egg cells in the ovary must be successfully fertilised by sperm cells from the pollen tube. The reproductive phase of plants, particularly male reproduction, is highly vulnerable to heat stress (Zinn et al., 2010; Müller and Rieu, 2016; De Storme and Geelen, 2014). Microsporogenesis, including the development of tetrads and unicellular microspores following meiotic divisions, is considered the most sensitive stage of pollen development to heat stress in plants (Sato et al., 2002; Endo et al., 2009; Begcy et al., 2019; Masoomi-Aladizgeh et al., 2020), with gametogenesis that follows possibly less heat sensitive, as reported in the binucleate stage of development in cotton (Masoomi-Aladizgeh et al., 2020). Molecular mechanisms that confer heat tolerance when the meiotic stage of pollen development is exposed to 40°C could be revealed by investigating related wild cotton species endemic to hot dry semi-deserts.

Wild species are valuable sources of novel genes with unique adaptive mechanisms that enable survival in inhospitable environments. Cotton (*Gossypium*) is an important crop that is commonly grown for natural fiber production across the world. The genus *Gossypium* includes eight groups of diploid species (n = 13) divided into multiple genomes (A-G, and K), of which C-genome species (*Gossypium robinsonii* and *G. sturtianum*) are endemic to Australia (Wendel and Cronn, 2003). Wild cotton species host many unique traits for resistance to abiotic stresses such as heat, drought and salinity (Mammadov et al., 2018) but after early work on the cytology and taxonomy of the genus, there are no reports on the mechanism of thermotolerance in these species. Hence, the applied goal of this research is to identify novel genes from distantly related species and introgress them into cultivated cotton without a yield penalty.

We sequenced genomic DNA from *G. robinsonii*, an Australian wild cotton C-genome species (2n = 2x = 26), and developed a draft genome for this species. This enabled us for the first time to use genome survey sequencing of a wild cotton (*G. robinsonii*), thereby translating predicted gene sequences for deep analysis of our *G. robinsonii* SWATH proteomic data. Mature pollen was collected for proteomic analysis after pollen cell lines of *G. robinsonii* had been exposed to moderate (36°C) and extreme heat (40°C). Observations that *G. robinsonii* plants set seed even at 40°C led to the hypothesis that heat stress during three stages of pollen development would elicit developmentally distinct proteomic patterns in mature pollen, and that while some proteins will reflect impaired metabolism as a result of heat stress, others play a direct role in heat stress tolerance.

## MATERIALS AND METHODS

### Plant materials

*Gossypium robinsonii* (2n = 2x = 26; Australian wild cotton) was grown in glasshouse conditions at 30/22°C day/night temperature, and 12/12 h light/dark photoperiod under natural light. The light intensity during the day cycle was maintained at a minimum of 600 µmol m^−2^ s^−1^ using supplemented light (Philips LED grow lights). Plants were fertilized weekly using Aquasol® soluble fertiliser (Yates, Australia) at the rate of 4 g per 5 l and watered daily. For Genome Survey Sequencing (GSS), young leaves grown under control conditions were collected. For proteomics, three stages of pollen development in *G. robinsonii* including tetrads (TE; 5 - 5.5 mm), uninucleate microspores (UN; 7 - 10 mm) and binucleate microspores (BN; 13 - 24 mm) were exposed to 36/25°C (moderate heat) or 40/30°C (extreme heat) for 5 d and corresponding mature pollen grain samples were collected and analysed in biological triplicate. Figure 1 shows the workflow of GSS on leaf and proteomics analysis of mature pollen heated at early stages of development.

**Figure 1.**
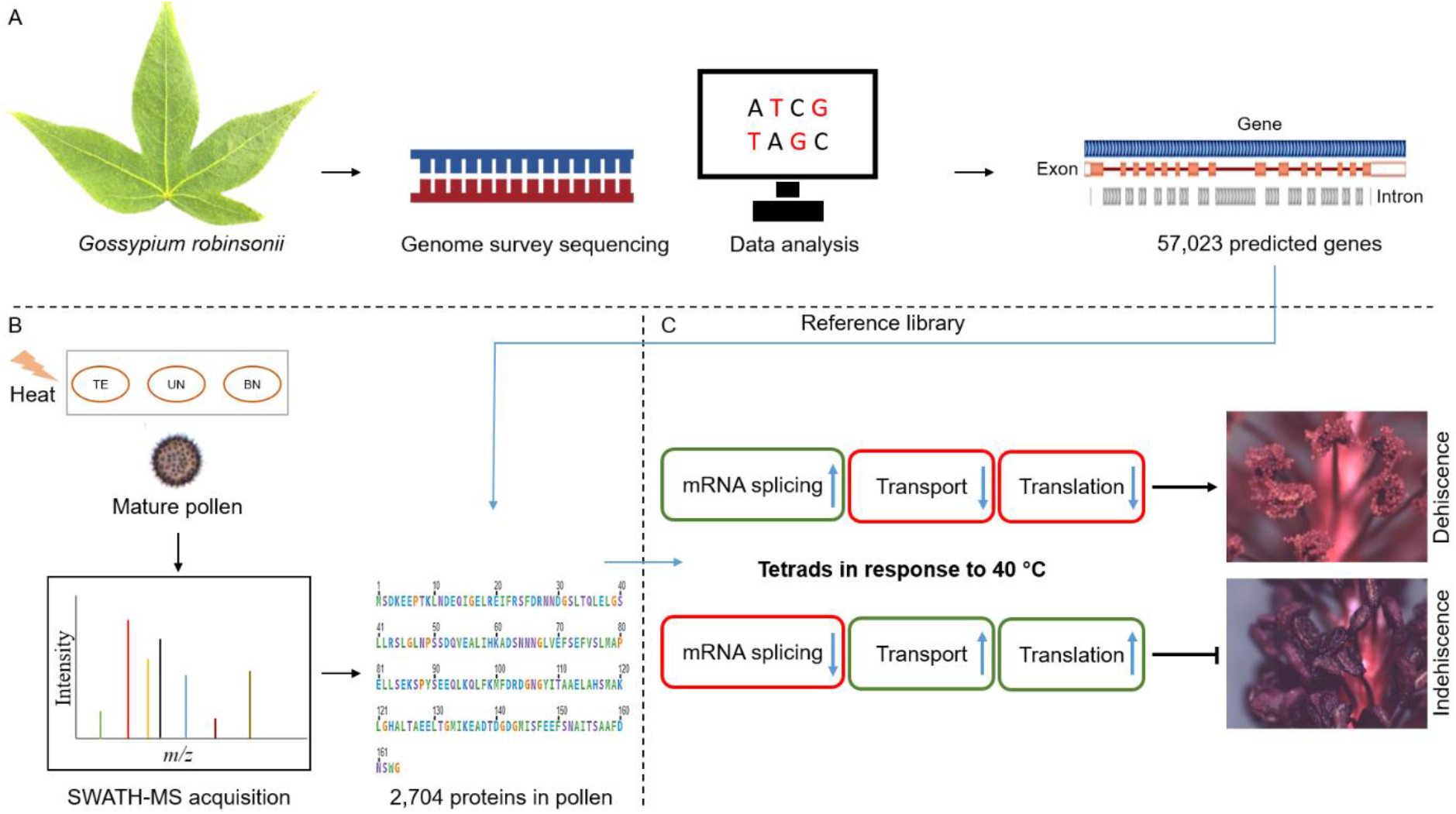
(A) Genome survey sequencing of *G. robinsonii* resulting in a total of 57,023 genes; (B) Exposure of three developmental stages of pollen development (TE, UN and BN) to 36 or 40°C for 5 d and proteomics analysis of the resulting mature pollen (p-TE, p-UN and p-BN); (C) The proposed thermotolerance mechanism of tetrads in *G. robinsonii*.

### DNA isolation

Total genomic DNA was extracted from approximately 100 mg young leaf tissue using a combination of a method described in Masoomi-Aladizgeh et al., (2016) and a DNeasy Plant Mini Kit (Qiagen, USA). Briefly, a lysis buffer for DNA extraction was prepared as described in Masoomi-Aladizgeh et al. (2016). The lysis buffer (1200 µl) in addition to βME (50 µl) were added to the ground leaf and samples were incubated at 65 °C for 10 min. Chloroform (600 µl) was added to each tube, and the mixture was vortexed vigorously, followed by centrifugation at 14,000 rpm at room temperature for 10 min. The supernatant (650 μl) was transferred into the QIAshredder Mini spin column to continue the isolation processes according to the manufacturer’s instructions. The isolated DNA was treated with 2 µl RNase, and the quality and quantity of the sample was assessed using agarose gel electrophoresis and a fluorometric procedure (Qubit Fluorometer, Invitrogen, USA) by the Beijing Genomics Institute (BGI, Shanghai, China).

### Whole genome sequencing

A paired-end library with insert size of 350 base pairs (bp) was constructed from randomly fragmented genomic DNA in the Beijing Genomics Institute (BGI). A BGISEQ-500 sequencer was utilized to generate sequencing data with read length of 150 bp. Quality of the reads was controlled and a stringent filtering process was carried out to obtain clean data for subsequent analyses as described by Li et al., (2010). These reads were deposited in the NCBI Sequence Read Archive (SRA) database and are available under Bioproject accession number PRJNA592601. The clean data were used for K-mer analysis to estimate the size of the genome, repetitive sequences and heterozygosis (Li et al., 2010a). The short-read sequences were assembled to contigs and scaffolds using SOAPdenovo (Xu et al., 2011; Li et al., 2010a). Five-kb non-overlapping sliding windows along the assembled sequence was used to calculate the guanine plus cytosine (GC) content and average sequencing depth among the windows.

### *Phylogenetic analysis of* G. robinsonii

The cotton internal transcribed spacer (ITS) regions (ITS1, 5.8S rRNA and ITS2) sequences were downloaded from the National Center for Biotechnology Information (NCBI) and compared with the ITS of *G. robinsonii* in this study. The alignment of these sequences was performed using ClustalW following default parameters, and a phylogenetic tree was constructed using MEGA X software with maximum likelihood inference. Data were analysed using a general time reversible (GTR) model with gamma distributed rate variation across five categories and invariant sites (Kumar et al., 2018). A total of 500 bootstrap replicates were run for this analysis.

### Gene prediction and annotation

For *de novo* gene prediction, scaffolds of larger than 1000 bp were processed in OmicsBox ver. 1.4.11. Repeat elements in the genome were identified and masked by searching the RepBase database (Bao et al., 2015) using RepeatMasker ver. 4.0.9 and Dfam ver. 3.0 (Smit et al., 2018; Hubley et al., 2016). Augustus was employed to predict genes from the repeat-masked genome using following parameters trained on *Arabidopsis thaliana*: Ab initio as gene finding method, Qmap threshold of 30, minimum read alignment of 11, minimum intron length of 32, minimum exon length of 300 and depth coverage of 20 (Hoff and Stanke, 2013). For gene annotation, BLASTx alignment was performed between the predicted genes and common databases including UniProtKB and Swiss-Prot (E-value < 1e−3). The predicted genes were also aligned to GO mapping (Götz et al., 2008) and InterProScan (Hunter et al., 2009) to obtain functional annotation.

### Protein isolation and peptide preparation

Proteins were extracted by a phenol-based protocol previously described by Masoomi-Aladizgeh et al. (2020). The concentration of proteins was measured by BCA protein assay (Pierce, Thermo Fisher Scientific, San Jose). A total of 85 μg proteins from each sample was digested in-solution with trypsin, which was then followed by desalting the resulting peptides using stage tips packed in house with SDB-RPS (3M, St. Paul, MO) as described in Hamzelou et al. (2020b). Peptide concentration was then measured using Micro BCA assay (Pierce, Thermo Fisher Scientific, San Jose) and an equal amount of peptide sample from each individual sample type was used for the subsequent analysis.

### Nano LC–MS/MS

To determine whether the draft genome of *G. robinsonii* assembled in this study would provide sufficient information for further proteomics analyses, an initial shotgun proteomic assessment was performed. Non fractionated peptides were analyzed by nanoflow LC–MS/MS (nanoLC-MS/MS) using a Q-Exactive, Orbitrap mass spectrometer (Thermo Fisher Scientific, San Jose, CA) coupled to an Easy nLC 1000 nanoflow liquid chromatography system as described in Hamzelou et al. (2020a).

### High pH (HpH) reversed-phase fractionation

For generation of an ion library, a small fraction of peptides from all samples was pooled and fractionated using a Pierce High pH Reversed-Phase Peptide Fractionation Kit (Pierce, Thermo Fisher Scientific, San Jose). Briefly, 60 µg of the pooled peptides was reconstituted in 300 μL of 0.1% TFA. The mixture was transferred into a high pH reversed-phase spin column and the flow-through fraction was retained after centrifugation at 3000 × g for 3 min. The same procedure was repeated with Milli-Q water to collect the wash fraction. Peptides were sequentially eluted using a total of eight gradient fractions (5%, 7.5%, 10%, 12.5%, 15%, 17.5%, 20% and 50%) of acetonitrile and triethylamine (0.1%). The peptide fractions were then vacuum-dried and resuspended in 24 µl of 0.1% FA.

### Spectral library creation and SWATH-MS acquisition

Samples were analyzed in two stages. First, in a data dependent mode for ion-library generation, followed by a data independent acquisition on SWATH mode, for peptide quantification (DDA and DIA, respectively). Both were performed using LC-MS/MS on a Triple TOF 6600 mass spectrometer (Sciex, USA) equipped with an Eksigent nanoLC 400 liquid chromatography system (Sciex, USA).

### Information dependent acquisition (IDA)

HpH fractionated peptides were injected onto a reversed-phase trap (Halo-C18, 160 Å, 2.7 µm, 150 µm x 3.5 cm) for pre-concentration and desalted with loading buffer. The peptide trap was then switched in line with the analytical column (Halo-C18, 160 Å, 2.7 µm, 200 µm × 20 cm). Peptides were eluted from the column using a linear solvent gradient of 5 - 35% of mobile phase B (0.1% formic acid, 99.9% acetonitrile) over 60 min at a flow rate of 600 nl/min. The reversed phase nano-LC eluent was subject to positive ion nano-flow electrospray analysis in an IDA mode. In IDA mode, a TOF-MS survey scan was acquired (m/z 350 - 1500, 0.25 s) with the 20 most intense multiply charged ions (2+ to 4+; exceeding 200 counts/s) in the survey scan being sequentially subjected to MS/MS analysis. MS/MS spectra were accumulated for 100 milliseconds in the mass range m/z 100 – 1800 using rolling collision energy.

### Data independent acquisition by Sequential Windowed Acquisition of all Theoretical Spectra (SWATH)

For SWATH-MS, peptides of the individual samples were separated over RP linear gradient of using the same LC and MS instruments as specified above with positive ion nanoflow electrospray mode. In SWATH mode, first a TOF-MS survey scan was acquired (m/z 350-1500, 0.05 s) then the 100 predefined m/z ranges were sequentially subjected to MS/MS analysis. A total of 100 variable windows were selected based on the intensity distribution of precursor m/z in IDA data. MS/MS spectra were accumulated for 40 milliseconds in the mass range m/z 350-1500 with rolling collision energy optimized for lower m/z in m/z window +10%. To minimize sample carryover, blank injections were performed between every sample injection. Additionally, sample data were acquired in a randomized order to avoid batch effect biases.

### LC–MS/MS data analysis

The raw data acquired on Q-Exactive were searched against two reference proteome files including the available *G. hirsutum* protein sequences in UniProtKB (79,242 entries, April 2021) and the protein sequences obtained from *G. robinsonii* in the present study (57,023 entries) using MaxQuant 1.6.17.0 for peptide to spectrum matching (Cox and Mann, 2008). The reference proteome of *G. robinsonii* was selected for subsequent proteomics analyses. For ion library generation, data in the IDA mode were processed in ProteinPilot ver 5.0.1 (Sciex, USA) using the Paragon algorithm with default parameters against the reference proteome of *G. robinsonii*. The obtained library was then imported into PeakView (version 2.2, Sciex, USA) as a reference ion library and matched against individual SWATH data files. The following matching criteria were applied: top six most intense fragment ions for each peptide, a maximum number of peptides of 100, 75 ppm mass tolerance, 99% peptide confidence threshold, 1% false discovery rate threshold and a 5-min retention time extraction window. The peak areas for peptides were extracted by summing the areas under curve (AUC) values of the corresponding fragment ions using PeakView. The summed peak areas of the peptides were normalized against the total abundance values of respective samples, then used for protein quantification. To assess differentially expressed proteins, comparison of protein abundances across respective sample groups was performed using two-sample student’s *t*-tests. Proteins with *p* < 0.05 and an expression fold-change ± 1.5 were considered significantly changed between treatments (Wu et al., 2016).

### Functional enrichment analysis

The differentially expressed proteins (DEPs) were functionally annotated in OmicsBox (ver. 1.4.1). Using Fisher’s Exact Test, the lists of up- and down-regulated proteins at each condition were compared with the annotated *G. robinsonii* proteome. Significantly enriched gene ontologies (GOs, *p* < 0.05) were selected for subsequent analysis. To demonstrate the tendency towards increased or decreased abundance under heat stress, the common enriched GOs were displayed in polar plots using GOplot (Walter et al., 2015).

## RESULTS

### Genome sequencing and assembly of short reads

To generate an initial draft genome of *G. robinsonii* (2n = 2x = 26, CC), DNA isolated from leaves was used for genome survey sequencing (GSS). A total of 133.53 Gb clean reads with sequence coverage of approximately 70-fold were generated from the small-insert (350 bp) library, after low-quality reads were filtered. These data were used for 17-mer analysis from which the estimated genome size was 1.91 Gb (Figure 2A; Table 1). Likewise, 17-mer analysis was used to calculate the heterozygous and repeat rates of the genome which were 0.36% and 84.2%, respectively.

**Figure 2.**
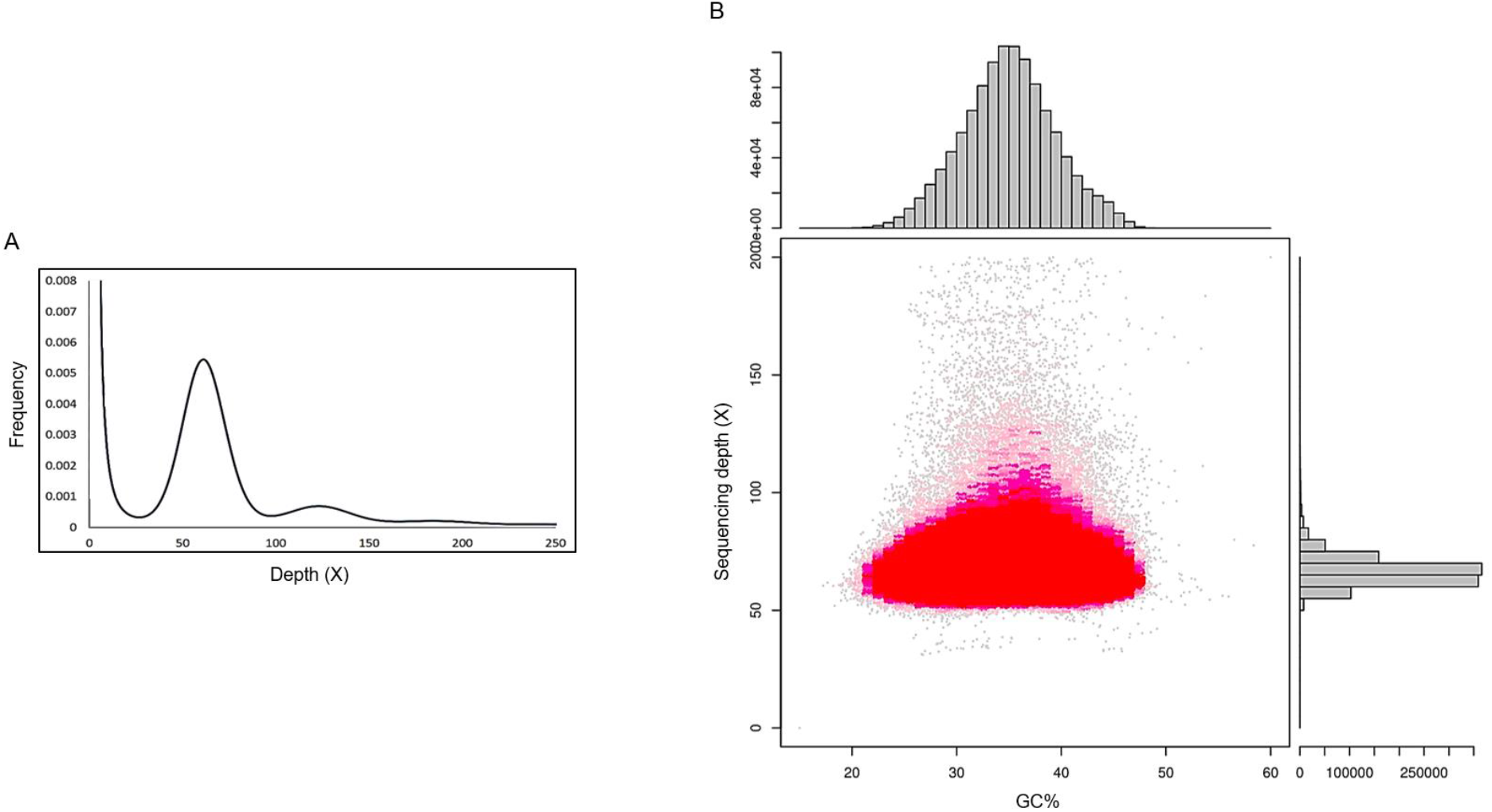
Genome characteristics of *Gossypium robinsonii*. (A) K-mer (K = 17) analysis to estimate the genome size of *Gossypium robinsonii* by the following formula: Genome size = K-mer number/peak depth. The *x*-axis is depth, whereas the *y*-axis indicates the proportion that represents the frequency at that depth divided by the total frequency of all depths; (B) The correlation analysis of guanine plus cytosine (GC) content and sequencing depth. The *x*-axis indicates the GC content, whereas the *y*-axis represents the sequence depth. The distribution of sequence depth is shown on the right side, while the distribution of GC content is at the top.

**Table 1.**
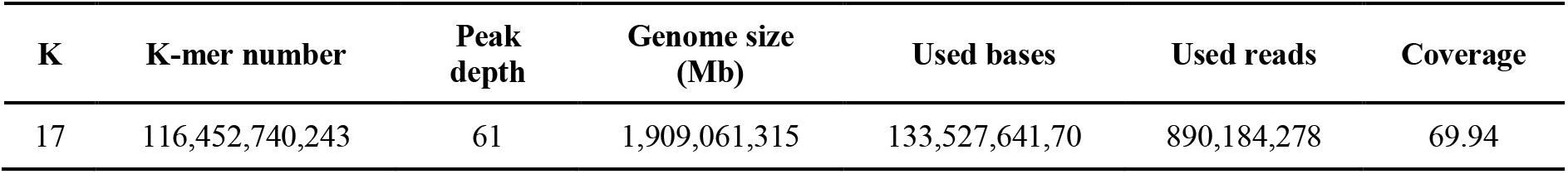
Statistical data from the 17-mer analysis

| K | K-mer number | Peak depth | Genome size (Mb) | Used bases | Used reads | Coverage |
| --- | --- | --- | --- | --- | --- | --- |
| 17 | 116,452,740,243 | 61 | 1,909,061,315 | 133,527,641,70 | 890,184,278 | 69.94 |

The clean reads were used as input to the SOAPdenovo program to create a *de novo* assembly. This assembly included a total of 50,428 contigs with an N50 of 6,837 bp. These contigs were subsequently assembled, resulting in 48,804 scaffolds with an N50 of 7,008 bp. Among all scaffolds, a total of 1,292,294 scaffolds were equal to or longer than 100 bp and 146,656 scaffolds were equal to or longer than 2,000 bp. The longest scaffold was 292,647 bp (Table 2). The average GC content of the *G. robinsonii* genome was 36.78% (Figure 2B).

**Table 2.**
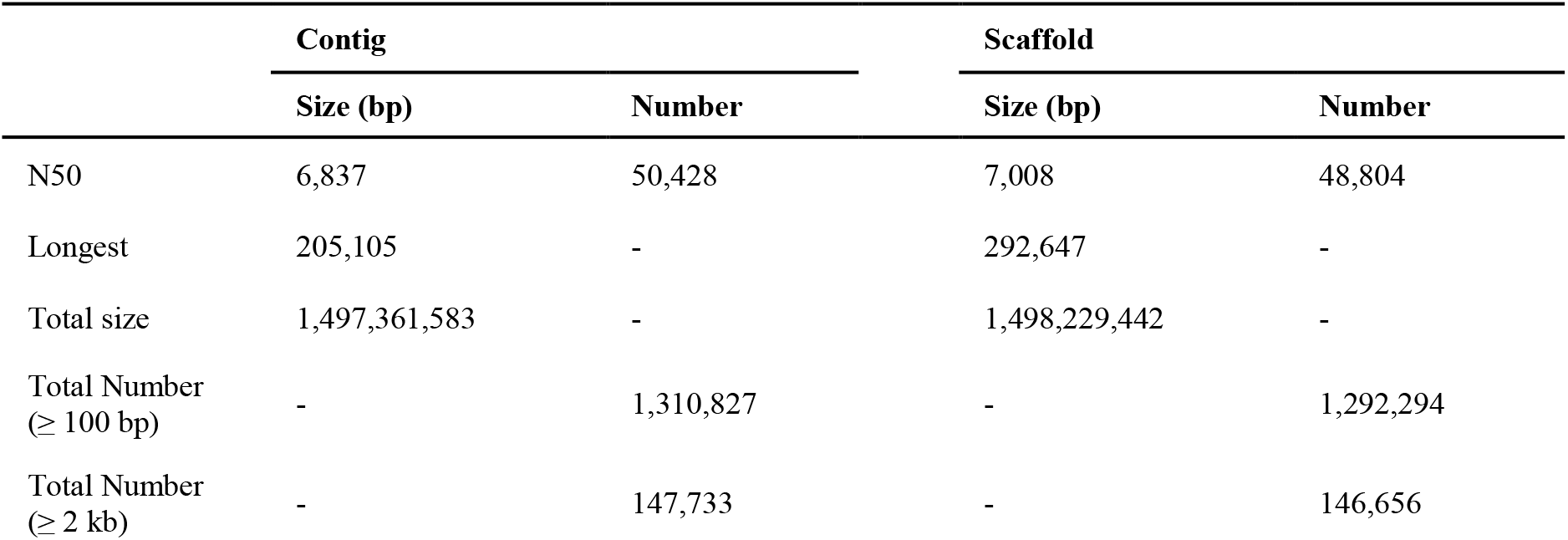
Statistics of *de novo* assembly results

### Repetitive elements analysis

A total of 1,292,294 scaffolds (≥ 100 bp) with an overall length of 1,498,229,442 bp were input into RepeatMasker to identify repeats in the assembled genome using Repbase. The results indicated that 54.8% of the assembled genome included repetitive sequences. The most abundant repetitive classes accounted for the long terminal repeat (LTR) elements consisting of 51.52% of the assembled genome, followed by DNA transposons (1.53%), simple repeats (1.06%), low complexity DNA sequences (0.37%), long interspersed nuclear elements (LINEs, 0.33%) and small RNA (0.03%). Gypsy/DIRS1 was the most common repeat among LTR elements, consisting of 47.99%, followed by Ty1/Copia (3.47%).

### *Confirmation of phylogenies in the genus* Gossypium

The *ITS* gene was used to establish phylogenetic clusters and species relationships for all 44 *Gossypium* species (Supplementary Data Table S1). Specifically, this enabled an assessment of the general reliability of the genomic data assembled above for *G. robinsonii* by establishing that it clustered among other Australian species that are believed to form a distinct taxonomic grouping. Cluster C grouped *G. robinsonii* with another C-genome Australian species (*G. sturtianum*) and the G-genome species, *G. australe, G. bickii* and *G. nelsonii*. The whole-genus phylogeny revealed six clusters (Figure 3), including cluster A that included 12 Australian species with the K-genome. Surprisingly, the *ITS* analysis suggested that *G. robinsonii* was more similar to the G-genome species in cluster C than its C-genome congener, *G. sturtianum*. Moreover, the substitution rate was higher in *G. robinsonii* compared with other Australian species, indicating a high rate of mutation over generations.

**Figure 3.**
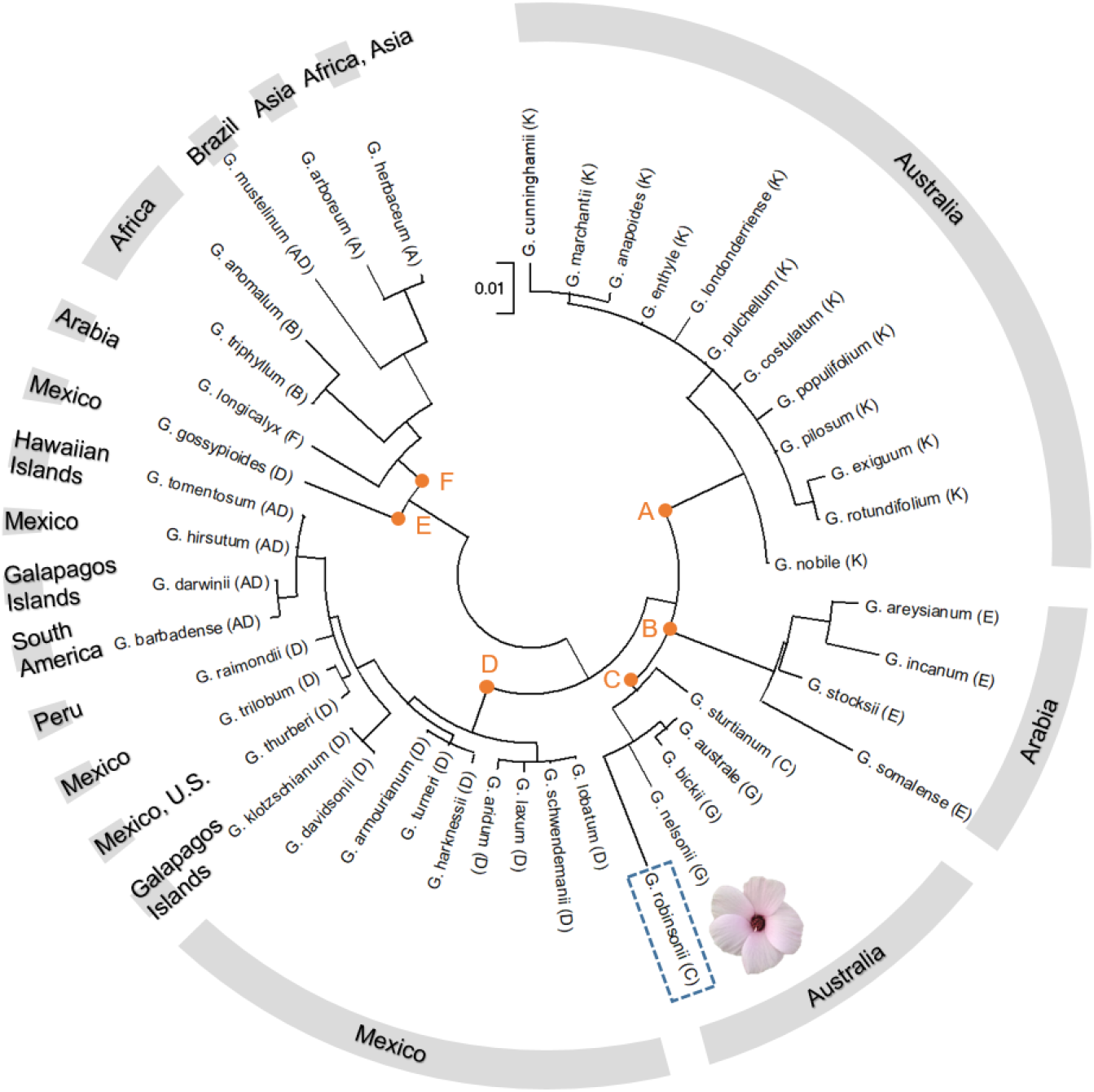
Phylogenetic relationship of *G. robinsonii* and other 43 cotton species using internal transcribed spacer (ITS) regions (ITS1, 5.8S rRNA and ITS2). The phylogenetic tree was constructed using MEGA X software with maximum likelihood inference. GTR model with gamma distributed rate variation across five categories and invariant sites was used for analyses (Kumar et al., 2018). The ITS sequences were downloaded from NCBI except *G. robinsonii* for which the gene was retrieved from the present study. The geographic location of the species was obtained from CottonGen database (http://www.cottongen.org). The branch lengths are drawn to scale with the bar, which indicates 0.01 nucleotide substitutions per site.

### Gene prediction and functional annotation

Based on the assembled genome of *G. robinsonii* with 242,945 scaffolds (> 1000 bp), we predicted a total of 57,023 genes which were assembled into a reference proteome sequence file. DNA fragments ranged from 102 to 21,615 bp in length with an average of 916.15 bp, all of which were retrieved from only 36,258 scaffolds. The putative genes were searched against UniProtKB and Swiss-Prot databases using BLASTx resulting in 30,847 (54%) functionally annotated genes.

### Extreme heat (40°C) during tetrad stage partially impaired dehiscence

The consequence of heat stress on reproductive success was investigated through monitoring dehiscence in p-TE, p-UN and p-BN. Dehiscence was successful at all stages of development when exposed to 36°C, and p-UN and p-BN underwent complete dehiscence in response to 40°C. Extreme heat, however, resulted in partial failure of dehiscence in p-TE. Of 17 squares heated at the tetrad stage, only 53% still underwent complete dehiscence while a further 18% partially dehisced and the remaining 29% were sterile.

### *Proteome profile of* G. robinsonii *for SWATH-MS analyses of pollen*

Prior to SWATH-MS analysis, the reference proteome sequences of *G. hirsutum* and *G. robinsonii* were employed for peptide to spectrum matching in trial experiments, resulting in identification of approximately 50% more unique peptides, from the same amount of starting material, when the *G. robinsonii* sequence was used. Hence, the *G. robinsonii* proteome was used for subsequent proteomics analyses. A proteome library was generated by pooling all samples analyzed, including p-TE36/40, p-UN36/40, p-BN36/40 and p-AN. Using the annotated *G. robinsonii* sequence as a reference, we obtained a total of 2,704 proteins from pollen (Supplementary Data Table S2). The functional annotations of these proteins, with associated biological process, cellular process and molecular function are shown in Figure 4 (only 30 top enriched categories). This SWATH ion library is available through the ProteomeXchange Consortium via the PRIDE partner repository (Perez-Riverol et al., 2019) with the dataset identifier PXD027097.

**Figure 4.**
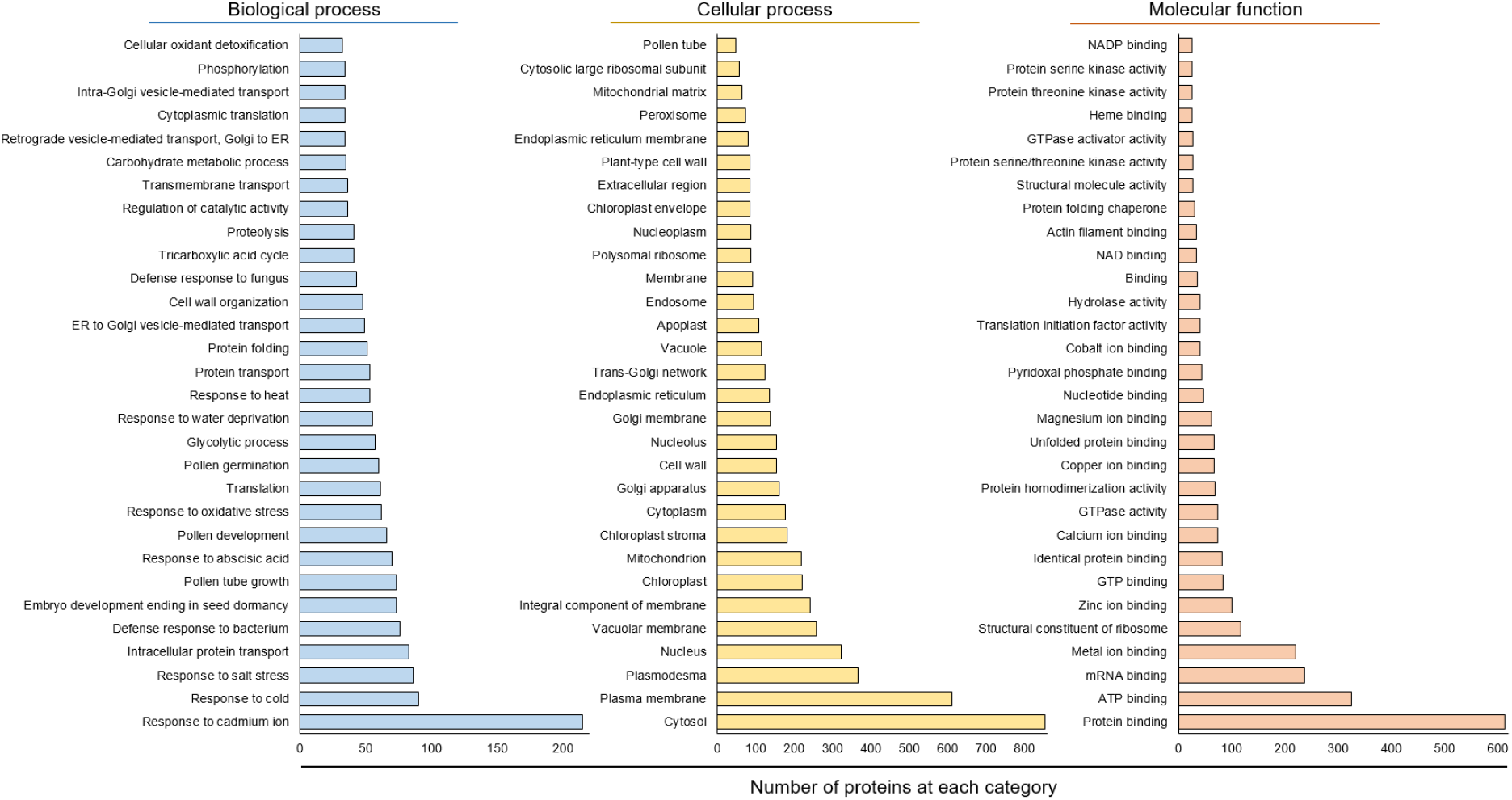
Functional Gene Ontology (GO) annotation of 2,704 proteins identified in pollen of *G. robinsonii* into 30 narrower categories that belong to ‘biological process’, ‘cellular process’ and ‘molecular function’. The *x*-axis shows the number of proteins, while the *y*-axis represents functional categories in which proteins are classified according to their abundance.

### Protein quantification of developing pollen heated at tetrad, uninucleate and binucleate stages

Using SWATH-MS for quantitative proteomic analysis, we identified 422, 489 and 94 DEPs in p-TE, p-UN and P-BN, respectively, after exposure to 36°C (moderate heat), compared with control pollen. Extreme heat (40°C) led to identification of only 297, 154 and 61 DEPs in p-TE40, p-UN40 and p-BN40, respectively (Supplementary Data Table S3), indicating a diminished response compared with the 36°C treatment and as above, a declining translational response when heat was imposed in late stages of pollen development. Differentially abundant proteins were visualised in an UpSet plot (Khan and Mathelier, 2017), highlighting that 196 DEPs were uniquely regulated in p-TE after exposure to extreme heat, whereas only 55 and 28 DEPs were unique to p-UN and p-BN, respectively, under the same conditions (Figure 5A and B). A total of 36 DEPs were common between the stages when plants were exposed to 36°C, while only 11 DEPs co-occurred in all stages under extreme heat. Significantly, we found that the number of DEPs in p-BN36 and p-BN40 decreased compared with p-TE and p-UN exposed to elevated temperature, in spite of all samples being derived from mature pollen (Figure 5C and D). Most notably, extreme heat caused changes in abundance of a large number of proteins in tetrads when compared with the later stages of development, which can be attributed to its distinctive response to high temperature and potentially its vulnerability to temperature stress.

**Figure 5.**
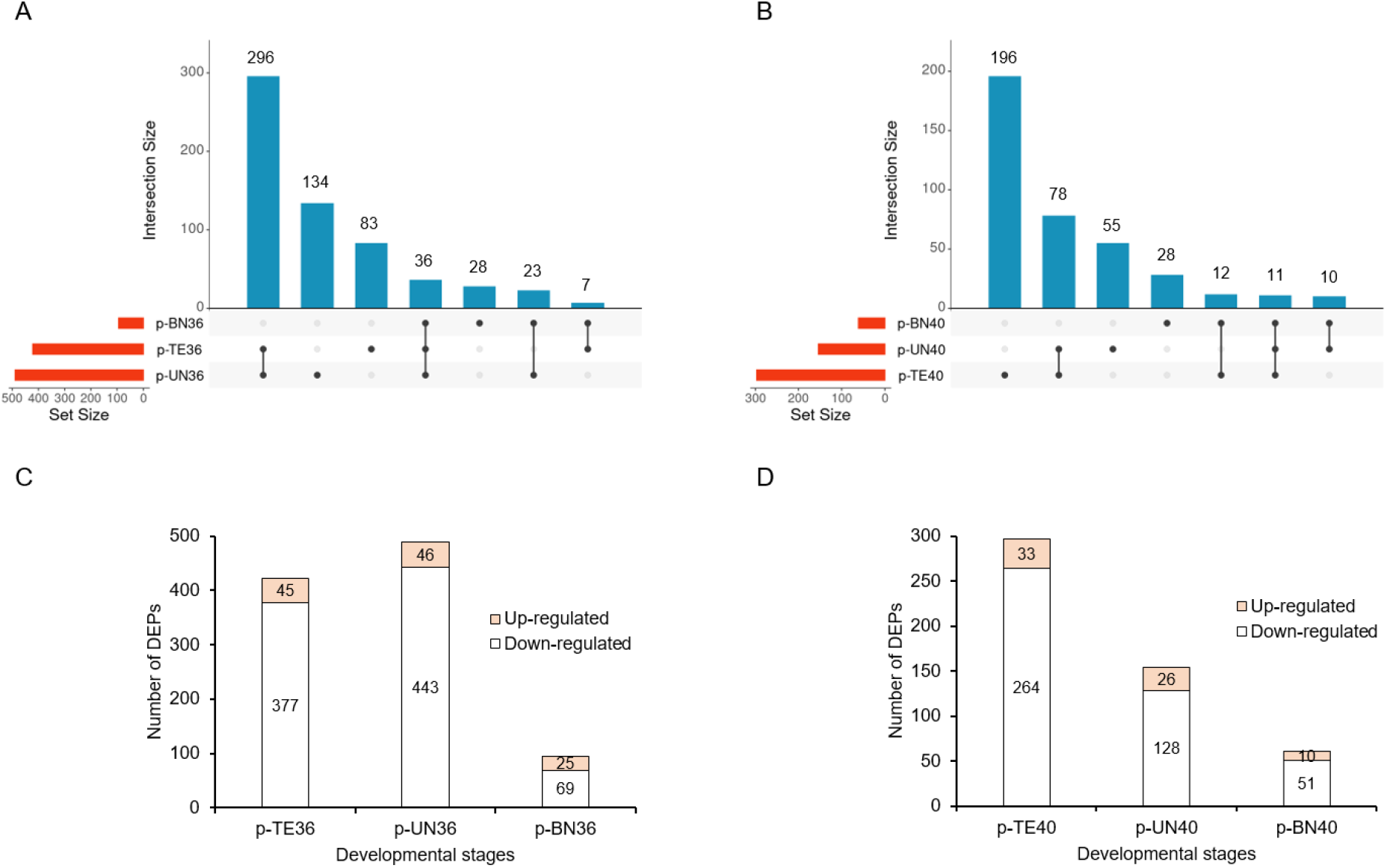
Differentially regulated proteins in mature pollen corresponding to tetrad cells (p-TE), uninucleate (p-UN) and binucleate (p-BN) microspores after exposure to 36 and 40°C for 5 d. (A and B) Upset plot illustrating the number of DEPs that are unique to each developmental stage (p-TE, p-UN and p-BN) after 5-d exposure to 36°C and 40°C, respectively; (C and D) number of up- and down-regulated proteins in p-TE, p-UN and p-BN stage when exposed to 36°C and 40°C, respectively.

### Functional annotation of p-TE, p-UN and p-BN in response to extreme heat

The top twenty most enriched gene ontology (GO) categories in biological process (Fisher’s exact test, *p*-value < 0.05) were plotted for DEPs under extreme heat (Figure 6). A detailed list of GO enrichment analysis is available in Supplementary Data Table S4. Enrichment analysis demonstrated that extreme heat caused an increased abundance of proteins involved in mRNA splicing and translation in p-TE40. Heat shock cognate 71 kDa protein (HSPA8/HSC70; Gr_Sca1204283G34) and glycine-rich RNA-binding protein RZ1A (RZ1A; Gr_Sca97552G39) were involved in mRNA splicing, whereas translation initiation factor 1A (Gr_Sca410536G29) and eukaryotic translation initiation factor 3 subunit D (Gr_Sca18748G18) were associated with cytoplasmic translation. All these key proteins increased in abundance after exposure of p-TE to 40°C. In addition, extreme heat increased the abundance of HSPA8/HSC70, 17.9 kDa class II heat shock protein (Gr_Sca284404G23) and cytosolic glyceraldehyde-3-phosphate dehydrogenase (Gr_Sca544237G31) in the response to hydrogen peroxide category in p-TE40 (Figure 6A). On the other hand, those functions mainly contributing to transport, such as proton export across plasma membrane, vesicle transport along microtubules, and Golgi to plasma membrane protein transport, were inhibited in response to extreme heat in p-TE. Importantly, Rab proteins (GTPases), which are key regulators of intracellular vesicle protein transport (Bhuin and Roy, 2014), decreased in abundance in p-TE40, indicating that intracellular transport might be down-regulated as part of the response of p-TE to extreme heat. These proteins included Ras-related protein Rab-2A (Gr_Sca778179G17 and Gr_Sca17313G6), Ras-related protein Rab-1B (Gr_Sca3776G55, Gr_Sca898430G34 and Gr_Sca790589G39) and Ras-related protein Rab-18 (Gr_Sca270469G20) (Figure 6B).

**Fig. 6.**
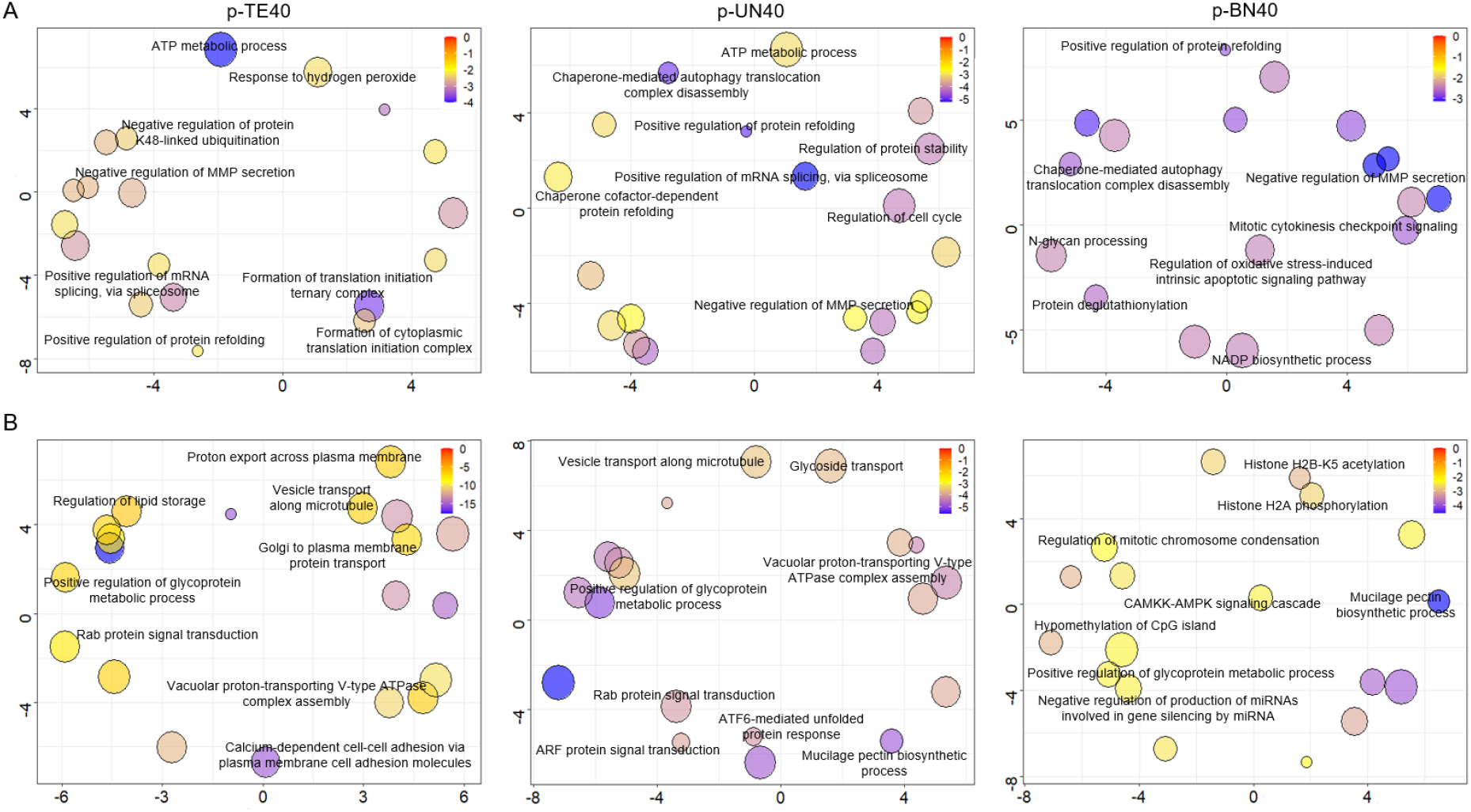
Most enriched categories (*p*-value < 0.05) in Biological Process up-regulated (A) and down-regulated (B) in p-TE, p-UN and p-BN in response to extreme heat. Each bubble represents a significantly enriched term in a two-dimensional space (Supek et al., 2015), with size that is proportional to the recurrence of the GO term in the whole Uniprot database. The scale indicates the log10 p-value, with red and blue representing the larger and smaller p-value, respectively. Of 20 most enriched GOs, only key categories were labelled in each plot.

In p-UN, extreme heat increased the abundance of two isoforms of heat shock cognate 71 kDa protein (Gr_Sca1204283G34 and Gr_Sca438049_G25) and glycine-rich RNA-binding protein RZ1A (Gr_Sca97552G39), which play a role in mRNA splicing and protein folding. Ankyrin-3 (Gr_Sca258696G41), 40S ribosomal protein S8 (Gr_Sca354645G21), 14-3-3 protein epsilon (Gr_Sca248462G23) and NEDD8-activating enzyme E1 catalytic subunit (Gr_Sca161791G12) also increased in abundance in p-UN40, all of which were involved in regulation of cell cycle (Figure 6A). As for p-TE40, proteins associated with transport were inhibited in p-UN after exposure to extreme heat. Rab proteins including Gr_Sca270469G20, Gr_Sca17313G6 and Gr_Sca790589G39 involved in transport were also down-regulated in p-UN40 to mitigate 40°C. Moreover, ADP-ribosylation factor 1 isoforms (Gr_Sca53462G15, Gr_Sca93217G3 and Gr_Sca18109G34), which are associated with ARF protein signal transduction and intracellular protein trafficking) decreased in abundance in p-UN40 (Figure 6B).

In p-BN, Gr_Sca438049G25 and glucosidase 2 subunit beta (Gr_Sca421233G26) were up-regulated in response to 40°C, both being involved in protein refolding. NADP dependent isocitrate dehydrogenase (Gr_Sca1122637G16), protein disulfide-isomerase (PDI, Gr_Sca49083G17) and Gr_Sca248462G23, which were enriched in NADP biosynthetic process, protein deglutathionylation and mitotic cytokinesis checkpoint signalling, respectively, also increased in abundance in p-BN40 (Figure 6A). Extreme heat, however, decreased the abundance of CAAX prenyl protease 1 homolog (Gr_Sca101458G58) which contributes to histone H2B-K5 acetylation, hypomethylation of CpG island, histone H2A phosphorylation and the CAMKK-AMPK signalling cascade. Additionally, proteins related to regulation of gene expression decreased in abundance in p-BN after exposure to extreme heat, including 60S ribosomal protein L5 (Gr_Sca687372G12), cold-inducible RNA-binding protein (Gr_Sca144232G10), SRSF protein kinase 1 (Gr_Sca381564G50), Gr_Sca101458G58 and programmed cell death protein 5 (Gr_Sca326285G39) (Figure 6B).

### p-TE notably decreased the abundance of proteins enriched in protein transport

Commonly enriched GOs across the pollen developmental stages (p-TE, p-UN and p-BN) were plotted to highlight the effect of extreme heat on the expression pattern of each category (Figure 7). The data indicated a notable decrease in abundance in mRNA binding (GO: 0003729), plant-type cell wall (GO: 0009505) and plasmodesma (GO: 0009506), after exposure of TE and UN to 40°C. Importantly, all proteins belonging to p-TE40 enriched in these categories decreased in abundance, indicating the importance of reduced protein expression in responding to extreme heat in the vulnerable stage of pollen development. Moreover, spliceosome complex (GO: 0005681), protein folding chaperone (GO: 0044183), positive regulation of mRNA splicing, via spliceosome (GO: 0048026), chaperone complex (GO: 0101031) and messenger ribonucleoprotein complex (GO: 1990124) increased in all stages in response to heat stress (Figure 7).

**Fig. 7.**
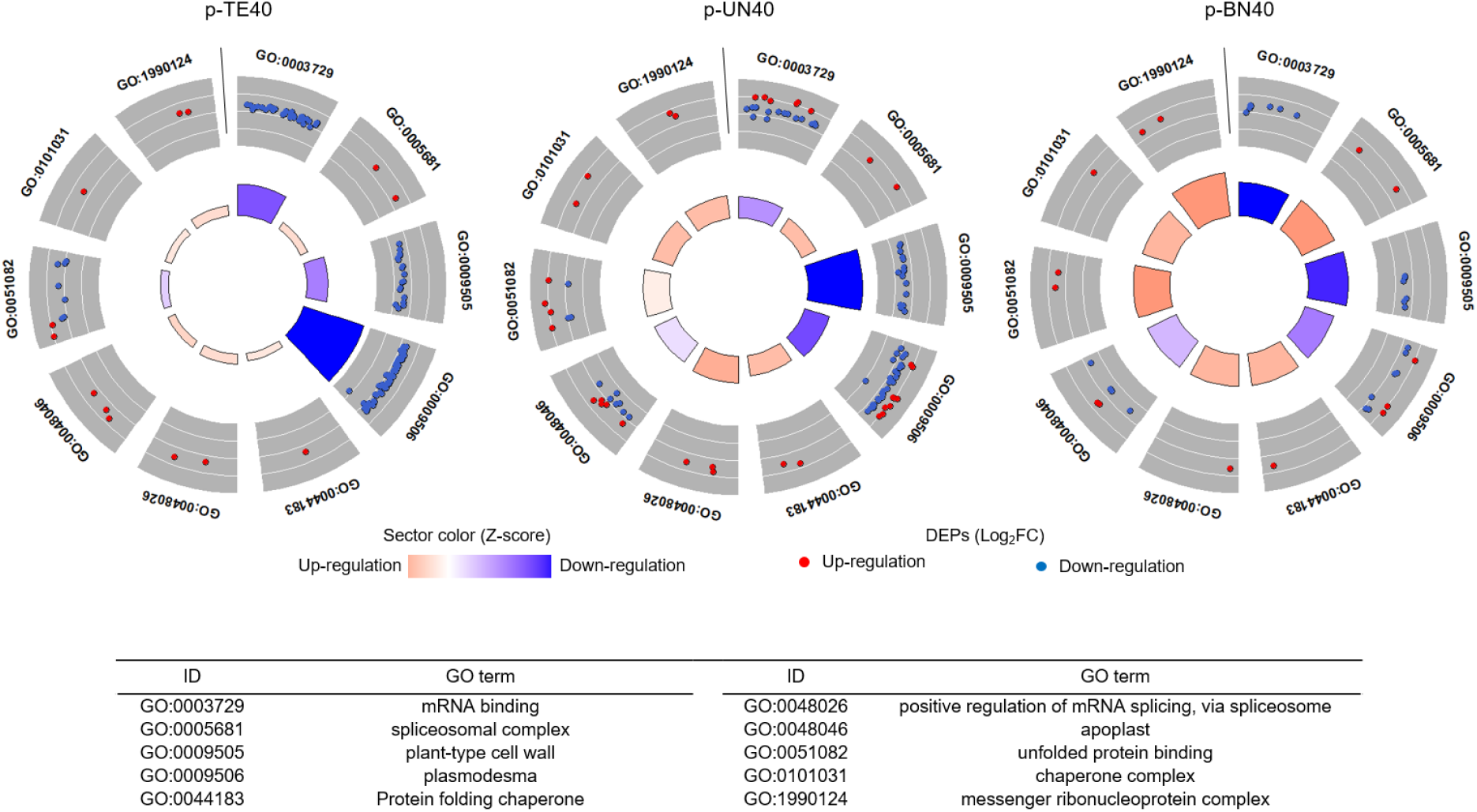
Expression patterns of ten commonly enriched GOs (*p*-value < 0.05) shared among three stages of pollen development (p-TE, p-UN and p-BN) after exposure to 40°C for 5 d. The outer ring shows the log 2-fold change of up- and down-regulated proteins enriched in a specific GO category. The sector size in the inner ring is proportional to the statistical significance (adjusted *p-*value) of each GO category, whereas the colour shows the tendency of the GO to increase or decrease in abundance in response to extreme heat according to its Z score, following this formula: 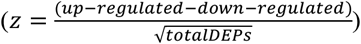.

## DISCUSSION

Heat stress during male gametophyte development results in a dramatically reduced fruit and grain production across a wide range of crops (Jagadish, 2020). Our recent investigation of a commercially cultivated cotton (Sicot 71) revealed that the meiotic stage of pollen development (tetrad cells) is extremely vulnerable to 40°C (Masoomi-Aladizgeh et al., 2020). We also found that up-regulation of non-essential pathways might contribute to the sensitivity of tetrad cells to heat stress in cotton. By inference, the progress of tetrad cells to dehiscent pollen would be promoted if most pathways were down-regulated and only essential proteins were actively translated as an adaptive response to extreme heat (Masoomi-Aladizgeh et al., 2021). With this hypothesis in mind, we turned to *G. robinsonii*, a wild cotton species from the arid zone of Western Australia, to discover whether this heat-adapted wild relative has evolved protein expression patterns during meiosis (tetrad cells) that are qualitatively distinct from what is seen in commercial cotton exposed to heat.

### Higher mRNA splicing in mature pollen corresponding to tetrads exposed to 40°C

A higher need for expression of heat-responsive genes has been reported when plants are exposed to abiotic stress; however, defects in mRNA splicing of these genes may occur due to insufficient splicing machinery (Cui and Xiong, 2015). Moreover, increasing evidence signifies the importance of splicing regulators as key plant stress-mediators and suggests that any defects in splicing factors would impair plant response to environmental fluctuations (Laloum et al., 2018). Defective splicing machinery could be the reason for accumulation of unprocessed HSP70 in maize pollen in response to 45°C, as reported previously (Hopf et al., 1992). Masoomi-Aladizgeh et al. (2021) also attributed the sensitivity of tetrad cells to potential defects in splicing machinery after exposure of cotton pollen to 38°C. In wild cotton (*G. robinsonii*), however, functional analysis demonstrated that the abundance of proteins associated with mRNA splicing increased in response to 40°C. This may be ascribed to the thermotolerance of tetrads in wild cotton species, enabling the development of viable mature pollen at daytime temperatures that render commercial cotton flowers sterile. HSPA8/HSC70, a constitutively expressed chaperone protein, is known to be associated with many cellular processes, under normal and stress conditions (Stricher et al., 2013; Liu et al., 2012). Glycine-rich RNA-binding proteins in Arabidopsis are involved in negative autoregulation at the post-transcriptional level, by which mRNA containing premature stop codons are degraded via the nonsense-mediated mRNA decay (NMD) pathway (Schöning et al., 2008). The up-regulation of RZ1A in *G. robinsonii* may be also involved in this pathway to improve splicing efficacy to mitigate heat stress. Proteins such as HSPA8 and RZ1A which we have identified in wild cotton are potential candidates for breeding better thermotolerance in cultivated cotton.

### Lower translation in mature pollen corresponding to tetrads exposed to 40°C

Downshift of global protein synthesis is an adaptive response to external stimuli, notwithstanding the strategic expression of key genes required for acclimation. The majority of differentially regulated proteins was down-regulated in p-TE (89%) in response to 40°C, indicating that the repression of translation plays a part in thermotolerance. A qualitatively different pattern of translation was reported by Masoomi-Aladizgeh et al. (2021) in cultivated cotton where translation was *higher* in response to 38°C in tetrads rather than being suppressed. The use of the wild cotton relative indicates that a strategic response to heat is required for thermotolerance, with only key genes translated upon stress. For instance, we identified Eif1a and Eif3d as up-regulated in p-TE40, both of which are associated with translation initiation. Eif1a is involved in ‘ribosome scanning’ for initiating translation, and is reported to increase tolerance to NaCl and osmotic stress in plants (Yang et al., 2017; Rausell et al., 2003). Eif3d also plays a key role in stress-induced translation and is involved in a mechanism underlying selective translation of essential genes required for cell survival (Lamper et al., 2020). Our finding is supported by Peredo and Cardon (2020), who highlighted that desiccation-tolerant algae massively down-regulated cellular metabolism, unlike the sensitive taxon, suggesting an energy-saving strategy under stress. Likewise, a study on grape (*Vitis vinifera*) indicated that the abundance of proteins involved in metabolic process decreased in response to 42°C (George et al., 2015). Repression of general protein synthesis in the meiotic stage of pollen development under extreme heat could lead to its thermotolerance, providing that translation of essential genes is maintained.

### Transport decreased substantially in mature pollen resulting from tetrads exposed to 40°C

Protein transport was reduced substantially when p-TE was exposed to extreme heat, which may be linked to adaptive mechanisms in tetrad cells of wild cotton that confer thermotolerance. In our previous study on cultivated cotton, we found that proteins associated with plasmodesma (GO:0009506) was highly up-regulated in the tetrad cells in response to 38°C, implying a high level of cell-cell communication that might lead to consumption of scarce resources, and consequently the vulnerability of tetrad cells (Masoomi-Aladizgeh et al., 2021). In sharp contrast, *G. robinsonii* suppressed cell-cell communication in p-TE40, demonstrated by down-regulation of proteins associated with plasmodesma. Rab proteins regulating intercellular transport were the key proteins down-regulated in p-TE40. This family of proteins is associated with a wide range of roles in plants, such as intracellular trafficking and responses to abiotic stress (Tripathy et al., 2021; Nielsen, 2020). For instance, it has been reported that mutations in NtRab2 blocked the localization of Golgi-resident, plasmalemma and secreted proteins, and also inhibited pollen tube growth in tobacco plants (Cheung et al., 2002). This agrees with our suggestion that the down-regulation of Rab-2A, Rab-1B and Rab-18 led to inhibition of intercellular transport in p-TE40. It appears that transport inhibition is required for tetrads of cotton to tolerate high temperature, and that Rab proteins could be potential tools in biotechnological approaches to improve heat tolerance.

## CONCLUSION

In this study, we generated an initial draft genome of *G. robinsonii*, an Australian wild cotton, and predicted a total of 57,023 genes which were assembled into a reference proteome sequence file. We found that GSS analysis of this obscure wild relative enabled us to identify 50% more unique peptides than would be achieved using sequences obtained from the cultivated species. Proteomic analysis of the wild cotton indicated that the thermotolerance of tetrads may be related to repression of translation of non-essential pathways, which resulted in decreased abundance of proteins in response to 40°C. Highly specific functions such as splicing were maintained as a survival mechanism. In addition, *G. robinsonii* suppressed cell-cell communication, which may conserve scarce energy resources that can be directed to essential processes throughout development. The application of mechanisms that are unique to this arid-zone wild species to cultivated cotton could increase the pollen thermotolerance, particularly through genetic intervention in meiotic processes.

## ACKNOWLEDGMENTS

F.M.-A. acknowledges support from Macquarie University in the form of iMQRES and MQRES scholarships. F.M.-A. acknowledges support from BioBam team (OmicsBox) in the form of a student scholarship for functional analysis of genes and proteins. Support from the Joyce W Vickery Scientific Research Fund from the Linnean Society of NSW and the Val Williams Scholarship in Botany from the Australian Plants Society (NSW) was seminal in initiating DNA sequencing and analysis. We appreciate use of the Plant Growth Facility (PGF), Australian Proteome Analysis Facility (APAF) and Macquarie University Faculty of Science and Engineering Microscope Facility (MQFoSE MF) and their valuable support. We also acknowledge the assistance of Yasmin Asar with phylogenetic analysis.

## AUTHORS’ CONTRIBUTION

F.M.-A. and B.J.A. conceived the original research plan. F.M.-A. cultivated and treated the plants, collected samples, performed the genomics and proteomics experiments, interpreted the results and prepared the original draft of the manuscript. K.S.K. assisted with SWATH-MS and proteomic data analysis. P.A.H and B.J.A. contributed to data interpretation and preparation of the final version of manuscript. All contributed to revision of the manuscript.

## DATA AVAILABILITY STATEMENT

The proteomics data were deposited in PRIDE (https://www.ebi.ac.uk/pride) under the accession number PXD027097. The genome sequence reads obtained by BGISEQ-500 are available in the NCBI Sequence Read Archive (SRA) at https://www.ncbi.nlm.nih.gov/sra. The Bioproject accession number is PRJNA592601 and the Biosample accession number is SAMN17125411.

## Notes

### Competing Interest Statement

The authors have declared no competing interest.

